# Directed evolution of split APEX peroxidase

**DOI:** 10.1101/452888

**Authors:** Yisu Han, Jeffrey D. Martell, Tess C. Branon, Daniela Boassa, David M. Shechner, Mark H. Ellisman, Alice Y. Ting

## Abstract

APEX is an engineered peroxidase that catalyzes the oxidation of a wide range of substrates, facilitating its use in a variety of applications, from subcellular staining for electron microscopy to proximity biotinylation for spatial proteomics and transcriptomics. To further advance the capabilities of APEX, we used directed evolution to engineer a split APEX tool (sAPEX). Twenty rounds of FACS-based selections from yeast-displayed fragment libraries, using three different yeast display configurations, produced a 200-amino acid N-terminal fragment (with 9 mutations relative to APEX2) called “AP” and a 50-amino acid C-terminal fragment called “EX”. AP and EX fragments were each inactive on their own but reconstituted to give peroxidase activity when driven together by a molecular interaction. We demonstrate sAPEX reconstitution in the mammalian cytosol, on engineered RNA motifs within telomerase noncoding RNA, and at mitochondria-endoplasmic reticulum contact sites.

## Introduction

APEX2 is an engineered variant of soybean ascorbate peroxidase that, unlike the commonly used horseradish peroxidase (HRP), retains activity when expressed in the cytosol, mitochondria, and other reducing environments within the cell^1,2^. This feature of APEX2, in addition to its versatile ability to catalyze the H_2_O_2_-dependent one-electron oxidation of a wide variety of small molecule substrates, has led to its widespread use for a variety of applications, including proteomic mapping of organelles^2–6^, proximity labeling of protein interactomes^7–9^, spatial mapping of cellular RNA^10^, electron microscopy^1,11–16^, H_2_O_2_ sensing^17^, and protein topology determination^1,2,16^.

APEX2 is typically fused to a protein or peptide to target it to a subcellular region or macromolecular complex of interest. For instance, we have targeted APEX2 to the outer mitochondrial membrane (OMM) and the endoplasmic reticulum membrane (ERM) of mammalian cells by fusing the APEX2 gene to transmembrane domains of proteins native to these subcellular locations^4,16^. These constructs were used for both EM^16^ and proteomic analysis^4^ of the OMM and ERM. While this APEX2 fusion strategy has enabled the study of many cellular regions and organelles, there are numerous compartments and structures to which APEX cannot be selectively targeted. For example, there is great interest in the biology of organelle-organelle contact sites, such as the junctions between mitochondria and ER, which participate in calcium signaling^18,19^, lipid synthesis^20–23^, and mitochondrial fission^24,25^. Yet all candidate protein fusions that could potentially target APEX2 to these contact sites, such as to the proteins Drp1^24^, Mfn2^26–28^, SYNJ2BP1^4^, and PDZD8^29^, would also target the peroxidase to locations outside of mito-ER contacts, such as throughout the cytosol^30^, along the cytoskeleton^31^, or over the entire OMM^4^.

Another application for which the APEX2 genetic fusion strategy may be unsuitable is profiling the interactomes of specific cellular RNAs. While several robust methods can identify RNAs that interact with specific proteins of interest^32–34^, the converse problem—identifying proteins that interact with a particular RNAs—is much more challenging using existing methods. One could envision fusing APEX2 to a high-affinity RNA-binding protein (RBP; for example, the bacteriophage MS2 coat protein^35^), allowing the peroxidase to be ectopically targeted to transcripts that are tagged with that RBP’s cognate RNA motif. However, a major concern would be the excess pool of catalytically active APEX2-RBP fusion protein that is not docked to the tagged RNA and can therefore produce off-target labeling that masks the specific signal.

A general solution to both these, and related, problems could be a split form of APEX2, in which two inactive fragments of APEX2 reconstitute to give an active peroxidase only when they are physically co-localized (**Figure 1A**). One could use an intersectional approach to restrict APEX2 activity specifically to sites of interest–such as mito-ER contacts, or specific RNA binding sites – thus eliminating the background labeling from off-target peroxidase activity.

**Figure 1.**
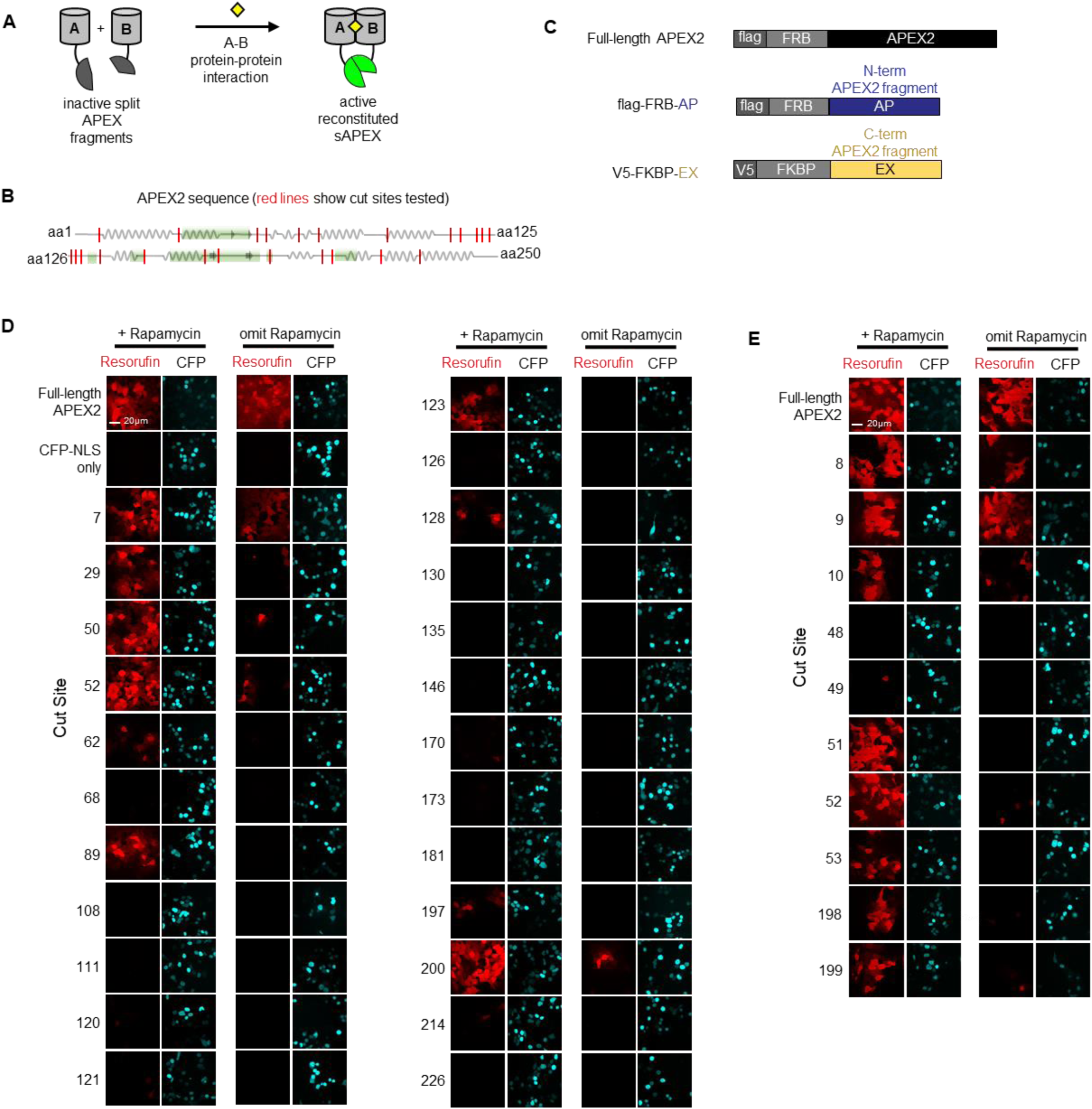
Split APEX design and screening of potential sAPEX cut sites. (**A**) Schematic overview of split APEX (sAPEX). Two inactive fragments (grey) can reconstitute to give active peroxidase (green) when driven together by a protein-protein interaction (PPI). The yellow square represents a chemical that can induce dimerization (**B**) The first screen tested 24 different cut sites. Their locations in the APEX2 protein sequence are indicated by the red vertical lines. Squiggles denote alpha helices. Grey arrows denote beta sheets. Areas shaded green are part of the heme-binding pocket. (**C**) N- and C-terminal sAPEX fragments selected for testing were fused to FRB and FKBP, respectively. (**D**) Initial screen of cut sites; split occurs after the indicated amino acid. For instance, cut site 7 splits APEX2 between residues 7 and 8. Pairs of constructs were introduced into HEK 293T cells by transient transfection, along with a CFP-NLS (nuclear localization signal) co-transfection marker. Cells were either treated with rapamycin for 24 h (left) or remained untreated (right). Subsequently Amplex UltraRed, a fluorogenic small-molecule peroxidase substrate, and H_2_O_2_ were added for 25 minutes, after which cells were fixed and imaged. Resorufin is the fluorescent product of Amplex UltraRed oxidation and indicates peroxidase activity. Scale bars, 20 µm. Three biological replicates were performed. (**E**) Second cut site screen, focused on residues surrounding T7, G50, and E200. Same assay as in (**D**). Two biological replicates were performed; representative images shown.

Although split protein reporters have been developed from green fluorescent protein^36,37^, HRP^38^, dihydrofolate reductase^39^, ubiquitin^40^, luciferase^41–43^, beta-galactosidase^44–47^, TEV protease^48^, Cas9^49–52^, and BioID^53^, splitting APEX2 presents new challenges. First, APEX2 requires a heme cofactor for its activity, and many cut sites would split the heme-binding pocket between the two fragments (**Figure 1B**). Second, in order for split APEX2 to be useful for a broad range of applications, the inactive fragments should have relatively low affinity for one another, such that reconstitution only occurs when the fragments are driven together by a molecular interaction. Not many known split proteins have low-affinity fragments, and it is challenging to engineer such a property in conjunction with high activity upon reconstitution.

To address the need for an interaction-dependent proximity labeling tool, we describe here the development of an evolved split APEX2 (sAPEX) system. Using a novel yeast display-based directed evolution approach that incorporates positive and negative selection steps, we have attempted to minimize interaction-independent association of the peptide fragments while maintaining high peroxidase activity upon reconstitution. Our resulting sAPEX fragment pair diverges substantially from its parental sequence and shows interaction-dependent reconstitution in multiple contexts in living mammalian cells. Our sAPEX tool adds to the proximity labeling toolkit, and in the future, it should extend the utility of APEX-based approaches to new areas of biology at higher spatiotemporal resolution.

## Results

### Structure-guided screening of potential APEX2 cut sites

We first sought to identify promising cut sites in the APEX2 enzyme using a chemically-inducible protein association system as a test platform. We selected 24 potential cut sites within solvent-exposed loops and turns between secondary structural elements (alpha helices and beta sheets) based on the crystal structure of wild-type ascorbate peroxidase^54^ (**Figure 1B**). Cut sites were also selected to avoid the creation of hydrophobic termini that might destabilize the fragments. All fragments were expected to lack a complete heme-binding pocket, with the exception of the large fragments produced from cut sites 7/8 and 29/30. We cloned each of the 24 fragment pairs as fusions to FKBP and FRB, whose interaction can be induced with the small molecule rapamycin (**Figure 1C**). We introduced the fragment pairs into HEK 293T mammalian cells using transient transfection and evaluated peroxidase activity in the presence or absence of rapamycin. Catalytic activity was detected using an established assay based on the membrane-permeable fluorogenic probe Amplex UltraRed, which is colorless but is converted into the red fluorophore resorufin upon peroxidase-catalyzed oxidization^55,56^.

Of the 24 fragment pairs tested, seven produced significant resorufin product, indicative of reconstituted activity (**Figure 1D**). For all cut sites except 7/8, much stronger activity was detected in the presence of rapamycin compared to when rapamycin was omitted, indicating that reconstitution was dependent on a protein-protein interaction. In the case of cut site 7/8, resorufin fluorescence was observed not only when rapamycin was omitted, but also when the large C-terminal fragment (amino acids 8-250) was expressed on its own (**Figure S1**), making this cut site unsuitable for our purposes. Cut sites within the heme binding cavity (130/131, 134/135, 146/147, 170/171, 173/174, and 181/182) produced fragments that completely failed to reconstitute activity (see **Figure 1B** and **Figure 2** for recolored full length APEX2 (PDB:10AG)).

**Figure 2.**
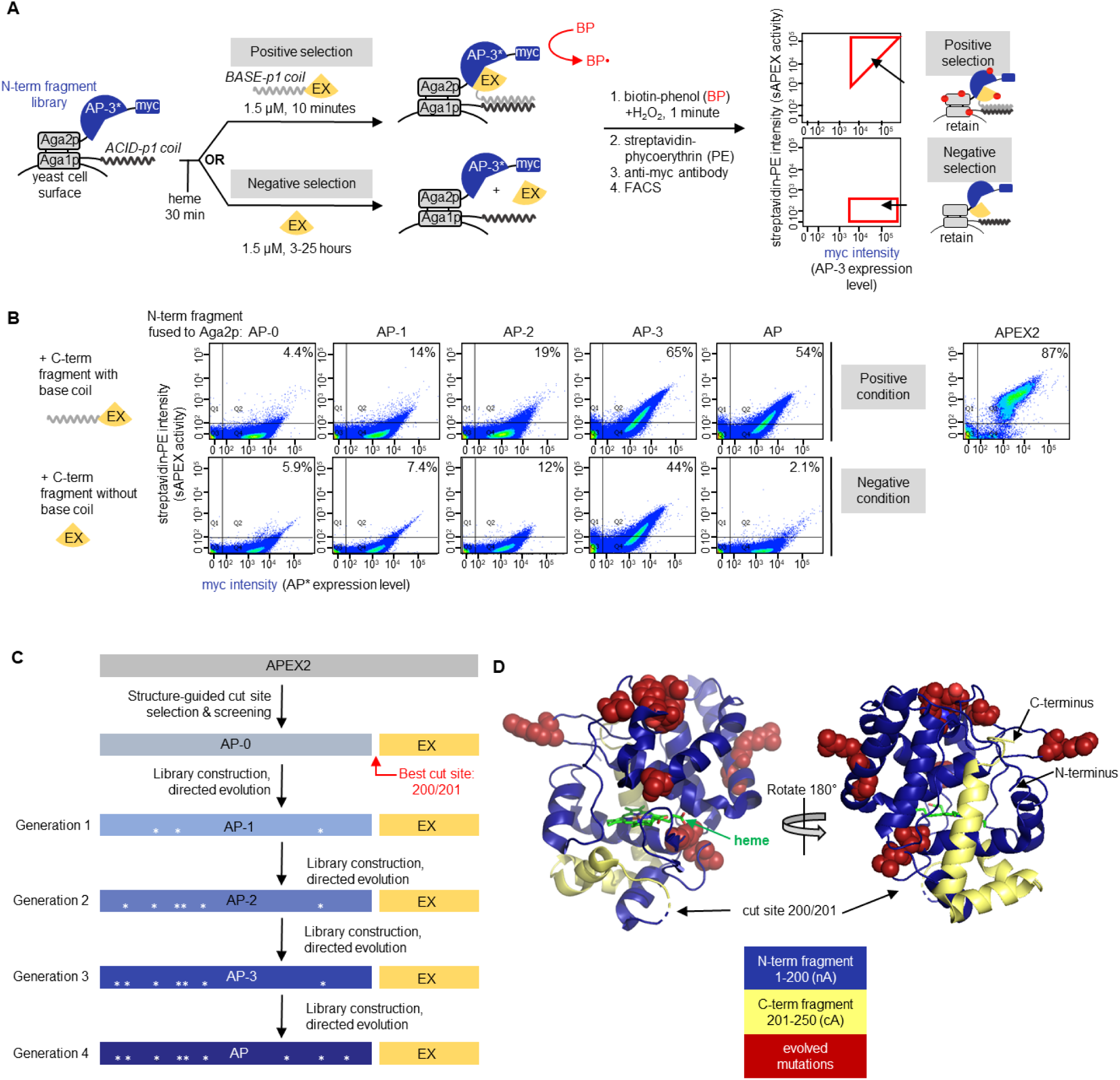
Yeast-display directed evolution and results. (**A**) Yeast display-based directed evolution scheme. The scheme shown here was used for Generation 4 selections, but the setup for other generations was similar (detailed in **Figures S3-S6**). A library of N-terminal fragments (“AP-3”, finalized clone from Generation 3 selections) was displayed on the yeast cell surface via fusion to the Aga2p mating protein. An acid coil was co-displayed, via fusion to Aga1p, to recruit base coil-fused C-terminal fragment (“EX”, amino acids 201-250 of APEX2). In the positive selection for high sAPEX activity, base coil-EX-GFP was incubated with the AP-3 yeast library for 10 minutes, then reconstituted peroxidase activity was detected by treating the cell mixture with biotin-phenol (BP) and H_2_O_2_. Cells with high peroxidase activity label themselves with biotin to a high extent^16^, enabling their enrichment via FACS after streptavidin-phycoerythryin (PE) staining. In the negative selection to deplete the sAPEX library of AP-3 fragments with excessively high affinity for EX, we incubated the AP-3 yeast library with EX-GFP protein lacking base coil for increasingly long time periods (see **Figure S6A**), then performed BP labeling. Cells with low streptavidin-PE signal were retained via FACS. Myc staining on the x-axis provides a readout of AP expression level. Note that the fluorescence from GFP was not measured during FACS; GFP was included to increase the solubility of EX. Hypothetical data are shown to illustrate the strategy for gate selection. (**B**) Summary of improvement of sAPEX activity and PPI-dependence throughout generations of selections in yeast. Full length APEX2 fusion to Aga2p was used as a benchmark for desired activity range. Yeast were prepared as in (**A**), with the indicated N-terminal fragment of sAPEX expressed on the yeast surface as a fusion to Aga2p. Purified C-terminal fragment (EX), either with or without acid coil, was added to the cells for 10 minutes or 7 hours, respectively. Then BP labeling, streptavidin-PE staining, and FACS were performed as in (**A**). AP-1, AP-2, and AP-3 are the first, second, and third generation N-terminal fragment clones, whose mutations are shown in (**C**). The percentage of myc-positive cells in the top right quadrant (Q2) is indicated in the top right corner of each FACS plot. Data are shown for one out of two biological replicates. (**C**) Summary of split APEX protein engineering. The name of the best N-terminal fragment clone to emerge from Generation 1 selection is “AP-1”, and so on, as indicated. The best clone to emerge from the Generation 4 selection is “AP.” Asterisks depict the locations of mutations within the protein sequence. (**D**) sAPEX split site and mutations in sAPEX relative to full-length APEX2^2^. The split site is between E200 and G201, and the N-terminal fragment (“AP”) and C-terminal fragment (“EX”) of sAPEX are colored blue and yellow, respectively. The nine residues mutated through directed evolution are colored red and rendered in space-filling mode; the original residues from the parent protein are depicted. Structures based on PBD ID: 1OAG^54^.

To more finely map the optimal cut sites, we performed a second round of screens, evaluating cut sites 1–3 residues away from promising sites identified in our initial screen (**Figures 1E and S1**). We ultimately identified three optimal cut sites—51/52, 89/90, and 200/201—all of which fell in solvent-exposed loops distal (>15 angstroms away) from the APEX2 active site and heme-binding pocket. For each cut site pair, controls in which either the N- or C-terminal fragments were expressed alone lacked any detectable peroxidase activity (**Figure S1**).

To assess the efficacy of our split APEX2 fragment pairs for applications in EM and proximity biotinylation, we evaluated their activity using the small molecule substrates diaminobenzidine (DAB)^1,16^ and biotin phenol (BP)^2–4^, respectively. Despite positive results in the highly sensitive Amplex UltraRed assay (**Figure 1D-E**), all three split APEX2 pairs produced far weaker DAB and BP staining than full-length APEX2 (**Figure 3B-D** and data not shown). For example, HEK 293T cells expressing full-length APEX2 exhibited very strong DAB staining after 15 minutes, while split APEX2 (cut site 200/201)-generated DAB stain under identical conditions was hardly detectable (**Figure 3B**). We found that the recently reported^57^ neighboring cut site, 201/202, exhibited activity similar to 200/201 (**Figure S2**).

**Figure 3.**
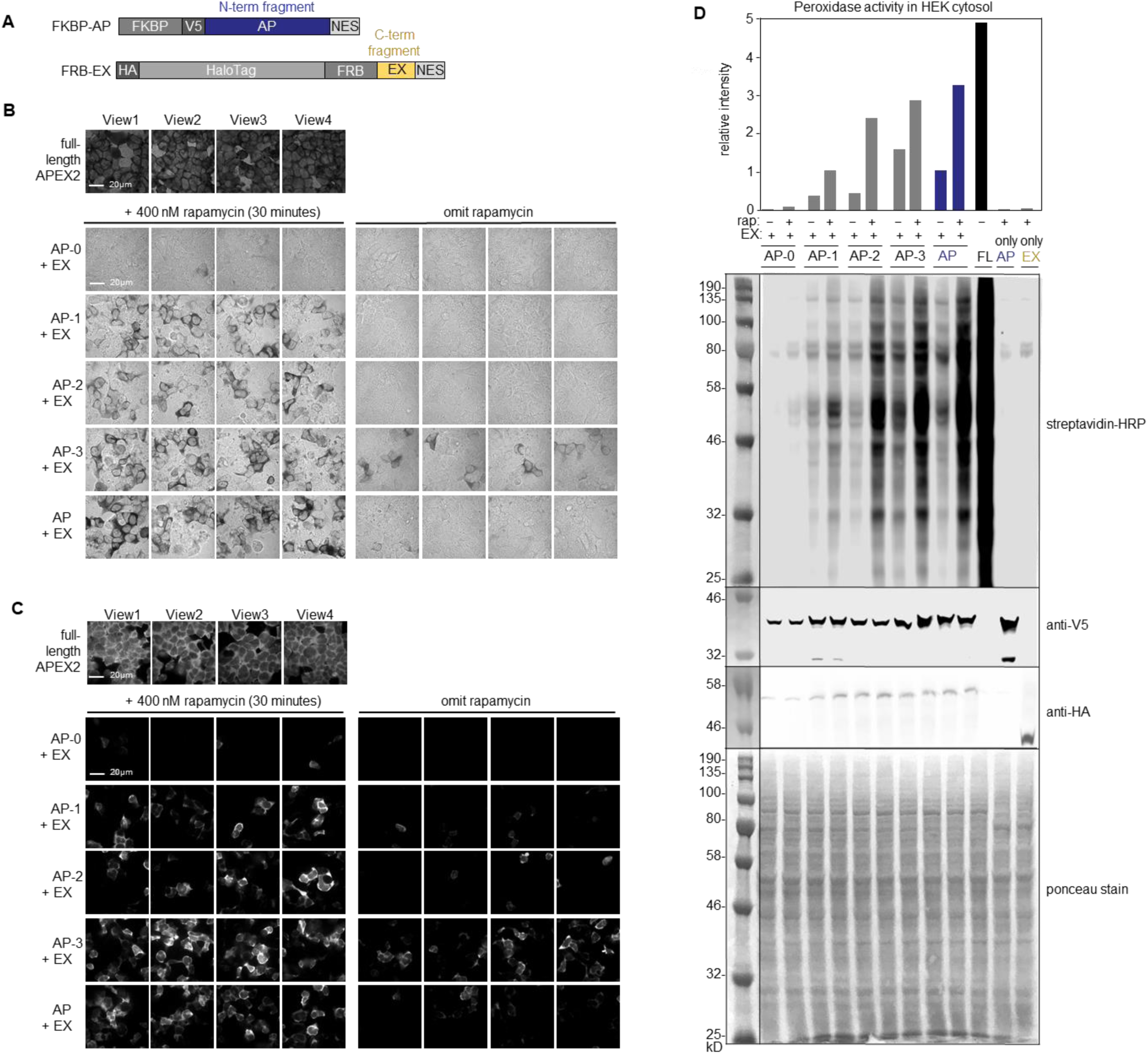
Comparing generations of evolved split APEX clones in mammalian cells. (**A**) Depiction of protein sequences for FKBP and FRB fusions to sAPEX for analysis in mammalian cells. N-terminal Halotag was fused to the N-terminus of FRB-EX to increase protein solubility. (**B**) Comparison of sAPEX variants in the mammalian cytosol, assayed by DAB (diaminobenzidine) polymerization activity. In the bright field images, dark regions indicate peroxidase activity. The indicated N-terminal variants of sAPEX were introduced by transient transfection into HEK 293T cells stably expressing FRB-EX, which were incubated with rapamycin for 30 minutes (left) or left untreated (right). We utilized HEK 293T cells stably expressing APEX2-NES as a benchmark, but because transfection and lentiviral transduction infection efficiencies are imperfect, this procedure resulted in a reduced number of comparable HEK 293T cells that would express both fragments. Cells were fixed and incubated with DAB and H_2_O_2_ for 15 minutes, as previously described^16^, to allow peroxidase-catalyzed polymerization of DAB. Four separate fields of view are shown per condition. Scale bar, 20 µm. Two biological replicates were performed. (**C**) Same assay as in (**B**) except FKBP-AP was introduced by lentiviral infection, and live BP labeling was used to detect peroxidase activity. Induced HEK 293T cells were treated with BP in the presence of H_2_O_2_ for 1 minute, then fixed and stained with neutravidin-AlexaFluor647 to visualize peroxidase-catalyzed promiscuous biotinylation^2^. Scale bar 20 µm. Four biological replicates performed. Additional fields of view shown in **Figure S7**. (**D**) Same assay as in (**C**) but with streptavidin blot readout. Two biological replicates performed. Quantitation of signal in each lane shown in bar graph at top. Anti-V5 and anti-HA blots detect expression of N-terminal and C-terminal fragments, respectively.

### A yeast display-based platform for split APEX2 evolution

The low activity of split APEX2 could be caused by a variety of factors, including poor stability, misfolding, aggregation of the fragments, or distorted geometry of the reconstituted active site and heme binding pocket, which could lead to low catalytic efficiency and/or poor heme recruitment. It is difficult to assess each of these potential problems, and even more difficult to fix them in a rational manner. Hence, we turned to directed evolution, which we and others have harnessed to improve or alter the properties of enzymes^16,38,58–62^. We selected yeast display-based directed evolution (**Figure 2A**) because, in contrast to other strategies such as high-throughput screening and phage display, yeast display allows processing of large mutant libraries (>10^7^) with large dynamic range – enrichment based on Fluorescence-Activated Cell Sorting (FACS) enables separation of highly active catalysts from moderately active ones, not just from inactive catalysts.

Our initial yeast display selection scheme exploited the yeast mating proteins Aga1p and Aga2p, which are displayed on the yeast cell surface and are joined by two disulfide bridges (**Figures 2A** and **S3A**). A library of yeast cells was generated, with each cell displaying on its surface a different mutant of the N-terminal split APEX2 fragment via fusion to the mating protein Aga2p. All cells concomitantly displayed the same C-terminal split APEX2 fragment as a fusion to Aga1p. The fragments were allowed to reconstitute for ~20 hours post-induction of protein expression. To read out the resulting peroxidase activity, APEX2-mediated biotinylation was initiated with biotin-phenol and H_2_O_2_, using standard conditions^16^. Yeast cells that display active reconstituted peroxidase under these conditions should promiscuously biotinylate proteins on their cell surface, which can be quantified using FACS (**Figures 2A** and **S3A**).

To establish this selection platform, we first created yeast Aga1p/Aga2p fusion constructs using the three promising split APEX2 pairs identified in the above screen. Because the 51/52 and 89/90 fragment pairs expressed poorly in yeast and gave no detectable activity on the yeast surface (*data not shown*), we proceeded with directed evolution of the 200/201 fragment pair. The C-terminal fragment (amino acids 201–250 of APEX2; henceforth called “EX”) was held constant while the N-terminal fragment (amino acids 1–200 of APEX2, called “AP-0”) was mutagenized using error prone PCR. Sequencing showed that our AP-0 library contained an average of 1.4 amino acid mutations per clone (**Methods**).

We performed four rounds of selection and observed that the activity of the yeast pool (measured by FACS) progressively increased (**Figure S3B**). We isolated 12 individual clones, characterized their activity on yeast using FACS and in the mammalian cytosol using microscopy, and combined mutations that appeared to be beneficial. The result was “AP-1”, which contains 3 mutations relative to the original APEX2. In a side-by-side comparison to the original split APEX2 fragment pair (AP-0 + EX) in HEK 293T cells, AP-1 shows noticeably improved activity in both DAB and BP labeling assays (**Figure 3B**).

To further improve the reconstituted activity of split APEX2, we performed another set of yeast selections in which the C-terminal EX fragment was not co-displayed on Aga1p but instead added as a purified, soluble protein (**Figure S4A**). This configuration allowed us to precisely control the concentration of EX added and the time of incubation, and to select for mutations that improved AP-1 expression and stability in the absence of EX co-translation. However, because the fragments only encountered each other on the yeast surface, where endogenous heme is not available, it was necessary to supply exogenous heme to the cells both during and prior to EX fragment addition. Four rounds of selection using this scheme produced the clone “AP-2”, which has three mutations beyond those in AP-1.

In a third-generation effort, to drive the fragments together using a protein-protein interaction (PPI) we again added EX as a soluble protein, but we used an artificially designed coiled-coil system, ACID-p1 and BASE-p1^63^, to recruit EX to the proximity of the N-terminal fragment (AP-2). This configuration mimics the split APEX2 reconstitution that would occur in our target biological applications. Four rounds of selection with gradual reduction of EX concentration and incubation time produced clone “AP-3”, which incorporates one additional mutation relative to “AP-2” and had noticeably higher activity than preceding clones in both DAB and BP labeling assays in HEK 293T cells (**Figure 3B-C** and **Figure S5**).

### FACS-based negative selections for reduced fragment affinity

An ideal split APEX2 fragment pair would have high catalytic activity when reconstituted, but low intrinsic binding affinity between the fragments, such that the fragments would only reconstitute when driven together by a PPI (**Figure 1A**). The clone AP-3, obtained after three generations of directed evolution, has much greater reconstituted activity (with EX) than does the original template (AP-0), but this activity was not dependent on PPI–induced co-proximation: using FRB and FKBP fusion proteins in HEK 293T cells, for example, we observed considerable DAB and BP signal in the absence of rapamycin (**Figure 3B-C**).

To preserve high reconstitution activity but decrease fragment affinity, we devised a new yeast display selection scheme that alternates between positive and negative selection rounds (**Figure 2A**). For the positive selection, we supplied the purified EX protein fused to the BASE-p1 helix that facilitates recruitment to Aga1p-ACID-p1. We performed BP labeling followed by streptavidin-phycoerythrin and anti-myc antibody staining. FACS was used to enrich cells with high SA/myc intensity ratios, as above. For the negative selection, we incubated the yeast with EX protein *lacking* BASE-p1 coil for extended periods of time (3 to 25 hours). AP-3 mutants that reconstituted with EX under these PPI-independent conditions were eliminated by FACS.

Starting with a library of 4.8 × 10^8^ AP-3 variants, we performed two rounds of positive selection and six rounds of negative selection (**Figure S6A**). By round 8, the yeast population was substantially depleted of cells that could reconstitute APEX2 activity upon addition of EX lacking the BASE-p1 coil (**Figure S6B**). We isolated 4 unique clones, characterized them by FACS, combined beneficial mutations, and re-tested in the mammalian cytosol. These experiments resulted in AP, the final split APEX2 (sAPEX) N-terminal fragment, which contains 9 mutations compared to the original APEX2 sequence. Mapping the positions of these 9 mutations onto the structure of wild-type ascorbate peroxidase, we observe that all lie in solvent-exposed regions, and none are at the interface between AP and EX (**Figure 2D**). Interestingly, half of the mutations are adjacent to cut sites we screened in **Figure 1D-E**, and many of the mutations were clustered in specific regions of the protein structure (**Figure 2D**).

### Characterization of enriched clones

Having completed 20 rounds of selection using three different yeast display configurations, we characterized key clones side-by-side. First, we prepared yeast displaying AP-0 (the starting template), AP (N-terminal fragment of final sAPEX), full-length APEX2, and AP-1 to AP-3 mutants, as fusions to Aga2p. Aga1p on these cells contained the ACID-p1 coil. We then supplied EX protein, either fused to (**Figure 2A**, *top row*) or lacking (*bottom row*) the BASE-p1 coil for proximity-dependent reconstitution. **Figure 2B** shows that the streptavidin-phycoerythrin (PE) staining (indicating reconstituted peroxidase activity) progressively increases from the template AP-0 to the finalized AP. However, the signal in the bottom row, reflecting proximity-independent reconstitution with EX (lacking the BASE-p1 coil), also increases, which is undesirable. After the implementation of negative selections in Generation 4, however, the untagged EX signal decreases for the AP clone. The EX+BASE-p1 coil signal remains high for AP, although not quite as high as that seen for AP-3. These observations on yeast indicate that the selections worked as desired and that our final clone AP combines the features of high reconstitution activity with low proximity-independent reconstitution.

We next tested whether these trends would hold in the mammalian cell cytosol. This environment is quite different from the yeast cell surface, as it is 37 °C instead of 30 °C, and a reducing rather than oxidizing environment. We also wished to test the sAPEX clones as soluble proteins rather than as membrane-anchored constructs with restricted geometry. Hence, we expressed the N- and C-terminal fragments from each stage of directed evolution as fusions to FKBP and FRB, respectively (**Figure 3A**). We first compared peroxidase activity with or without rapamycin using our DAB assay (relevant for EM applications^1,16^), which is much less sensitive than both Amplex UltraRed and BP assays, thus enabling us to rigorously compare the activity of our sAPEX fragment pairs (**Figure 3B**). The original sAPEX template, AP-0, gave barely detectable DAB staining, while AP-1 was dramatically improved, and AP-3 gave the strongest staining. However, as observed in yeast, AP-3 also gave significant signal in the absence of rapamycin, indicating high PPI-independent reconstitution. In contrast, the final AP + EX clones displayed high DAB staining in the presence of rapamycin, but nearly undetectable staining in the absence of rapamycin.

Analogous experiments using the BP assay—relevant for spatial proteomics^2,3^ and transcriptomics^64^—showed a similar trend (**Figure 3C** and **Figure S7**). While AP-3 showed high activity following rapamycin treatment, it also revealed background labeling in the – rapamycin condition. In contrast, the final sAPEX pair, AP + EX, displayed +rapamycin activity comparable to that of AP-3, but minimal – rapamycin activity. We also characterized BP labeling by lysing the cells and blotting the cell lysate with streptavidin-HRP (**Figure 3D**). The PPI-dependence of sAPEX makes it a versatile tool for cellular applications such as those outlined in the introduction.

### sAPEX reconstitution on a target RNA

As APEX-catalyzed proximity biotinylation has demonstrated utility for mapping protein interactomes^7–9^, there has been interest in using APEX2 to also map the interactomes of target nucleic acids – RNAs and individual genomic loci. In theory, this could be accomplished by fusing full-length APEX2 to programmable RNA- or DNA-binding proteins (such as CRISPR-Cas13d or Cas9, respectively)^65–67^ that specifically target the RNA or genomic locus under investigation. Alternatively, target RNAs or DNA loci could themselves be tagged with motifs that specifically bind APEX2 fusion proteins^68^. However, each of these approaches would be plagued by pools of excess catalytically active APEX2 that is unbound to the target of interest. This is exemplified by recent studies that used dCas9 to recruit APEX2 to specific genomic loci^66,67^, and a separate study that used the bacteriophage MS2 RNA coat protein (MCP) to recruit promiscuous biotin ligase variants to RNAs appended with the cognate MS2 RNA motif^68^. An analogous APEX2-MCP fusion system would also likely suffer from background biotinylation that overwhelms the specific signal from RNA-bound APEX-MCP.

We reasoned that our sAPEX system could potentially be used to alleviate this problem by fusing the evolved AP and EX fragments to orthogonal RNA-binding proteins, and selectively reconstituting APEX2 peroxidase activity on target RNAs that are appended with both of the cognate protein-binding RNA motifs (**Figure 4A-B**). As proof-of-principle, we sought to apply this strategy to TERC, the noncoding RNA component of the telomerase ribonucleoprotein (RNP), which synthesizes the ends of chromosomes in many clades of eukaryotes^69^. In addition to providing the template for telomere synthesis, the TERC ncRNA is also thought to serve as a structural “scaffold” onto which the other holoenzyme components assemble^70^. Critically, proper biogenesis of functional telomerase RNPs can be recapitulated when TERC RNA is transiently expressed from a plasmid, even if the RNA is appended with exogenous sequences at its 5′ end^71^. We therefore designed a series of variants in which the TERC 5′ terminus is appended by cassettes of motifs recognized by RNA-binding proteins. For this purpose, we chose to express AP and EX as fusions to the MS2 and PP7 bacteriophage nucleocapsid proteins—respectively termed MCP and PCP—which recognize disparate RNA motifs with exquisitely high affinity and specificity (cognate RNA K_D_ ~1nM, non-cognate RNA K_D_ ~1mM)^35,72,73,74^ (**Figure 4A-C**). Anticipating that sAPEX reconstitution might be sensitive to the specific geometry with which the AP and EX fragments are co-proximated, we designed a series of TERC variants that positioned MS2 and PP7 stem loops in different orientations (**Figure S8**).

**Figure 4.**
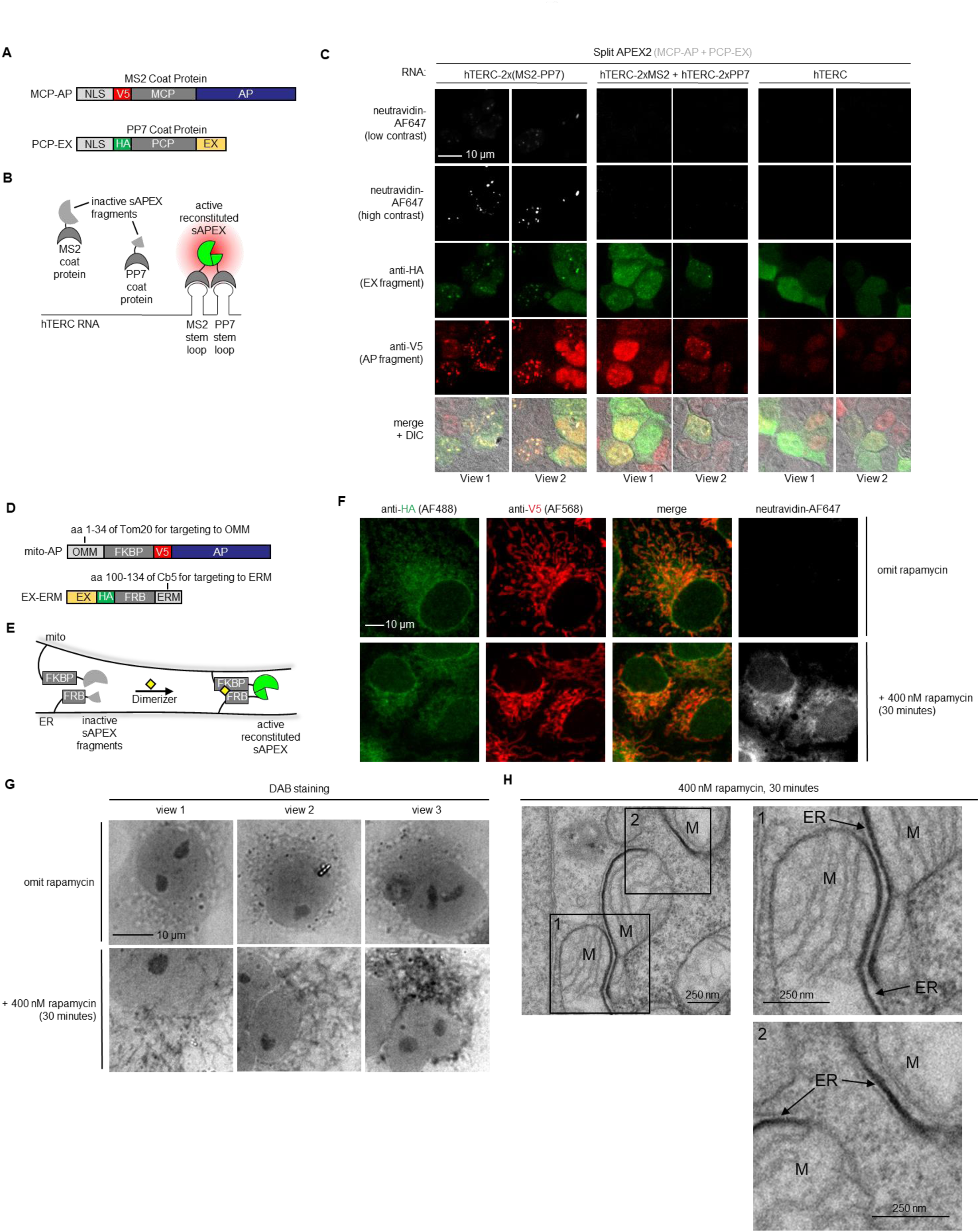
Testing split APEX on RNA binding sites and at mito-ER contacts. (**A**) Constructs used to test sAPEX targeting to RNA. MCP and PCP are the bacteriophage coat proteins that bind the MS2 and PP7 RNA stem-loops, respectively. Protein fusions are targeted to the nucleus by an N-terminal nuclear localization signal (NLS). (**B**) Schematic overview of sAPEX applied to interaction mapping of specific cellular RNAs. Cells expressing sAPEX (with N and C-terminal fragments fused to MCP and PCP, respectively) would localize peroxidase activity and labeling only to hTERC (human telomerase RNA component) RNA sites with adjoining MS2 and PP7 stem-loops. (**C**) Initial testing of sAPEX targeting to RNAs. HEK 293T cells stably expressing the MCP-AP construct shown in (**A**) were transfected with PCP-EX expression plasmid and the indicated RNA constructs. Twenty-two hours later, cells were subjected to *in situ* proximity biotinylation with BP and H_2_O_2_, fixed, and then stained with neutravidin-AlexaFluor647 to visualize reconstituted peroxidase activity, anti-HA antibody to visualize EX expression, and anti-V5 antibody to visualize AP expression. Two fields of view are shown per condition. Scale bar, 10 µm. This experiment has three biological replicates. Additional fields of view are shown in **Figure S9**. (**D**) Constructs used for targeting AP and EX fragments of sAPEX to the outer mitochondrial membrane (OMM) and ER membrane (ERM), respectively. (**E**) Schematic overview of sAPEX applied at mito-ER contacts. Inactive fragments (grey) fused to FKBP and FRB can reconstitute to give active peroxidase (green) when driven together by an inducible protein-protein interaction (PPI) with the addition of rapamycin (yellow diamond). (**F**) sAPEX reconstitution in COS7 mammalian cells. COS7 cells stably expressing EX-FRB-ERM from (**D**) were infected with lentivirus containing OMM-FKBP-AP. Forty-six hours later, cells were incubated with heme for 90 minutes prior to incubation with BP in heme-free media in which rapamycin was added or omitted, as indicated. BP labeling was initiated by the addition of H_2_O_2_, and after 1 minute, cells were fixed and stained with neutravidin-AlexaFluor647 to visualize reconstituted sAPEX activity. HA and V5 staining show the localizations of total EX and total AP, respectively. The top row is a negative control with rapamycin omitted. Scale bar, 10 µm. Additional fields of view are shown in **Figure S10**. This experiment has four biological replicates. (**G**) Reconstituted sAPEX has reactivity against DAB in COS7 mammalian cells. COS7 cells stably expressing EX-FRB-ERM from (**D**) were infected with lentivirus containing OMM-FKBP-AP. After 48 hours, cells were incubated with heme for 90 minutes prior to the rapamycin incubation in heme-free media for 0 or 30 minutes. Cells were then fixed, and DAB labeling was performed for 15 minutes. In the bright field images, dark regions indicate peroxidase activity. Scale bar, 10 µm. This experiment has three biological replicates. (**H**) sAPEX can be used as genetically-encoded reporter for EM. Samples from (**G**) with 30-minute rapamycin incubation were analyzed by EM. Dark staining from osmium tetroxide recruitment to sAPEX-generated DAB polymer is observed exclusively at sites where ER is in close proximity to mitochondria. Scale bars, 250 nm. This experiment represents a single biological replicate.

Expression of MCP–AP and PCP–EX resulted in biotinylation that was dependent on the presence of a TERC RNA bearing both of the cognate MS2 and PP7 motifs (**Figure 4C** and **Figure S9**). Critically, we observed no such biotinylation when individual MS2 and PP7 hairpins were localized on separate TERC RNA constructs. Moreover, the labeling pattern appeared punctate, as would be predicted from localizing peroxidase activity to the discrete subnuclear foci characteristic of interphase telomerase^75^. Our data suggest that sAPEX activity can be specifically reconstituted on a target RNA by nucleating protein fragment assembly on a structured RNA cassette.

### sAPEX reconstitution at mitochondria-ER contacts

In mammalian cells, an estimated 5–20% of the mitochondrial outer membrane makes intimate contact (<70 nm gap) with the membrane of the endoplasmic reticulum (ER)^19^. These mito-ER contacts are thought to be important for a variety of functions and signaling processes, from mitochondrial fission to lipid synthesis^18,19,20–23,24,25^. Recent work has identified proteins that reside at mito-ER contacts and may play a role in tethering the membranes together^4,29^. A major goal of this field is to comprehensively characterize the molecular composition of these contact sites to better understand how they mediate important cellular processes. To pave the way for such future efforts, we tested whether sAPEX activity could be reconstituted at mito-ER contact sites.

Mito-ER contacts are delicate and easily perturbed structures. Overexpression of various proteins, such as SYNJ2BP or even green fluorescent protein, can lead to dramatic distortion of one or both organellar membranes^4,76^. Because our optimized sAPEX fragment pair, AP + EX, does not exhibit PPI-independent reconstitution, we reasoned the tool may be suitable for reconstitution at mito-ER contacts without major perturbation of organellar structure. **Figures 4D-F and S10** show AP-FKBP targeted to the OMM and EX-FRB targeted to the ERM in COS7 cells, which are flat and thin, facilitating visualization of mitochondria and ER structures. We observed BP/streptavidin staining in cells treated with rapamycin for 30 minutes. Consistent with the PPI-dependence of sAPEX reconstitution, no BP labeling was observed when rapamycin was omitted. We also stained the COS7 cells for mitochondrial and ER markers and observed minimal disruption of organellar morphology, both before and after rapamycin addition (**Figure S10B-C**).

To examine sAPEX at mito-ER contacts at higher resolution, we examined mito-ER reconstituted sAPEX activity using electron microscopy (EM). We fixed the above COS7 cells, and overlaid with DAB and H_2_O_2_ to allow reconstituted sAPEX to catalyze the oxidative polymerization and local deposition of DAB^1^. Bright field imaging showed dark threads corresponding to stained mitochondria in the 30-minute rapamycin-treated samples, but not in the untreated samples (**Figure 4G**). We next stained the rapamycin-treated samples with OsO_4_ to deposit electron-dense osmium on the DAB polymer, then embedded and sectioned the samples for EM. EM imaging revealed a dark stain, corresponding to regions of reconstituted sAPEX activity, exclusively at contact sites between mitochondria and ER (**Figure 4H**). Two zoomed views show that DAB staining filled the ~25 nm gap between the OMM and ERM, but was absent from isolated ER and mitochondrial membranes. Notably, the mitochondrial and ER membranes were not grossly perturbed, as was observed in previous studies in which overexpressed reporters induced non-native mito-ER contacts and organelle aggregation^16^.

## Discussion

Using a combination of rational design and yeast-display directed evolution, we have engineered a low-affinity sAPEX protein complementation assay (PCA) with robust activity upon reconstitution driven by co-proximation. After three generations of evolution, we evolved a variant of split APEX2 with robust reconstitution, but which was prone to PPI-independent reconstitution. To overcome this limitation, our final directed evolution strategy implemented a negative selection that eliminated yeast clones with high fragment affinity. The final engineered sAPEX fragments, AP and EX, possess a total of nine mutations relative to the starting APEX2 template, clustered at solvent exposed regions, that collectively improve the PPI dependence of this system while maintaining high catalytic activity of the reconstituted form. This yeast-display platform could be extended to engineer other PCA systems. High-affinity fragment pairs such as those of split GFP result in spontaneous and irreversible reconstitution^77^; similarly, split HRP irreversibly reconstitutes in the ER without a PPI^38^. Split YFP utilizes fragments that reconstitute in a more PPI-dependent manner^78,79^, but it still suffers from background fluorescence, especially at high expression levels, and from irreversible YFP self-assembly^80,81^.

Given the purportedly broad array of novel RNAs and RNA-mediated processes that have eluded mechanistic dissection^82^ and the growing number of diseases now thought to be mediated by aberrant RNA-protein interactions^83,84^, there is a great need for identifying the interaction partners of specific RNAs. Conventional approaches address this problem by capturing and enriching a target RNA along with its bound interaction partners, either using affinity-tagged antisense oligonucleotides that isolate endogenous transcripts^85–88^, or by affinity tagging the transcript of interest itself^89,90^. However, these approaches are often confounded by the abundance of the target transcript, the unpredictable performance of RNA-based affinity tags^91^, and nonspecific RNA-protein interactions formed during lysis and enrichments^92^.

Using RNA scaffolding motifs to target the reconstitution of sAPEX eliminates high off-target background signal from protein overexpression. sAPEX demonstrates promise for BP labeling around specific RNAs, which indicates potential future applications to elucidate interactomes and interaction partners of an RNA of interest. Moreover, since the RNA domain we used to recruit AP and EX was designed to be functionally independent of the RNA to which it is appended, it should in theory be adaptable to other RNA targets^93^. A downside to this approach, however, is the need to express the transgenic, tagged RNAs that may not properly fold, localize, or function. As an alternative strategy, one might use RNA-targeting CRISPR-Cas systems to recruit AP and EX to endogenously expressed RNAs^65^. This approach may require substantial optimization to juxtapose the AP and EX fragments in the proper orientation, and may be influenced by the sequence, structure, or proteome of the targeted RNA.

The ability to reconstitute APEX2 activity at an organellar junction represents a promising step towards gaining greater understand of subcellular locales that were previously intractable to genetic targeting. We found that the reconstitution of sAPEX at mito-ER contact sites was PPI-dependent, despite multiple days of co-translation of the sAPEX fragments on the membranes of these organelles that are known to come into contact. Importantly, sAPEX did not greatly perturb the morphology of these organelles, as shown by fluorescence and electron microscopy. We note that some optimization was required to attain this PPI-dependence; for instance, we changed the promoter of OMM-FKBP-EX from CMV to human ubiquitin promoter (hUBC) to reduce the overall protein expression level, which reduced PPI-independent reconstitution. sAPEX potentially provides the ability to investigate and map the protein and RNA residents at other organelle-organelle contact sites beyond mito-ER junctions.

Unlike most other PCAs, sAPEX possesses the versatile capability to react with many different substrates, enabling its use for a wide array of applications. Because sAPEX activity can be reconstituted in highly specific subcellular regions that are intractable for single-gene constructs, it expands the toolkit of proximity labeling technologies.

## Acknowledgements

FACS was performed at the Koch Institute Flow Cytometry Core (MIT) and Stanford Shared FACS Facility. S. Han (Stanford) synthesized neutravidin AlexaFluor647. This work was supported by NIH grants to A.Y.T. (R01-CA186568), to D.B. (R01-GM086197), and to M.H.E. (P41-GM103412) for support of the National Center for Microscopy and Imaging Research. D.M.S. was supported by HHMI/Jane Coffin Childs Fellowship and by J. Rinn (U. C. Boulder). T.C.B. was supported by Dow Graduate Research and Lester Wolfe Fellowships.

---

**Figure S1.**
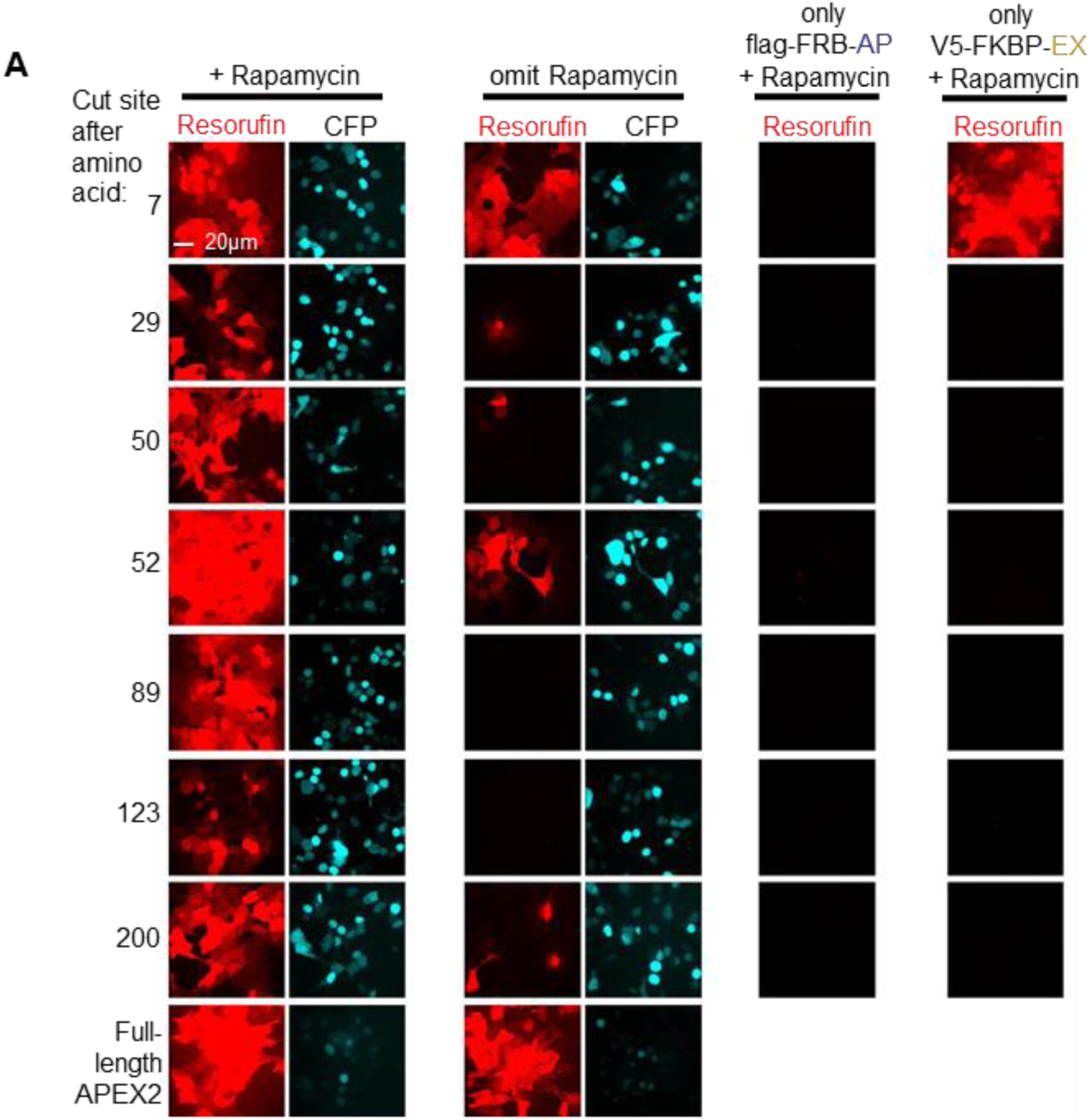
Screening of potential sAPEX cut sites. Pairs of constructs were introduced into HEK 293T cells by transient transfection, along with a CFP-NLS (nuclear localization signal) co-transfection marker. Here, for more stringent examination of promising cut sites, each fragment was transfected for individual expression in HEK 293T cells. Cells were either treated with rapamycin for 24 h (left column, and individual fragments) or left untreated (second column). Subsequently Amplex UltraRed, a fluorogenic small-molecule peroxidase substrate, and H_2_O_2_ were added for 25 minutes, after which cells were fixed and imaged. Resorufin is the fluorescent product of Amplex UltraRed oxidation and indicates peroxidase activity. Scale bars, 20 µm. The experiment has one biological replicate.

**Figure S2.**
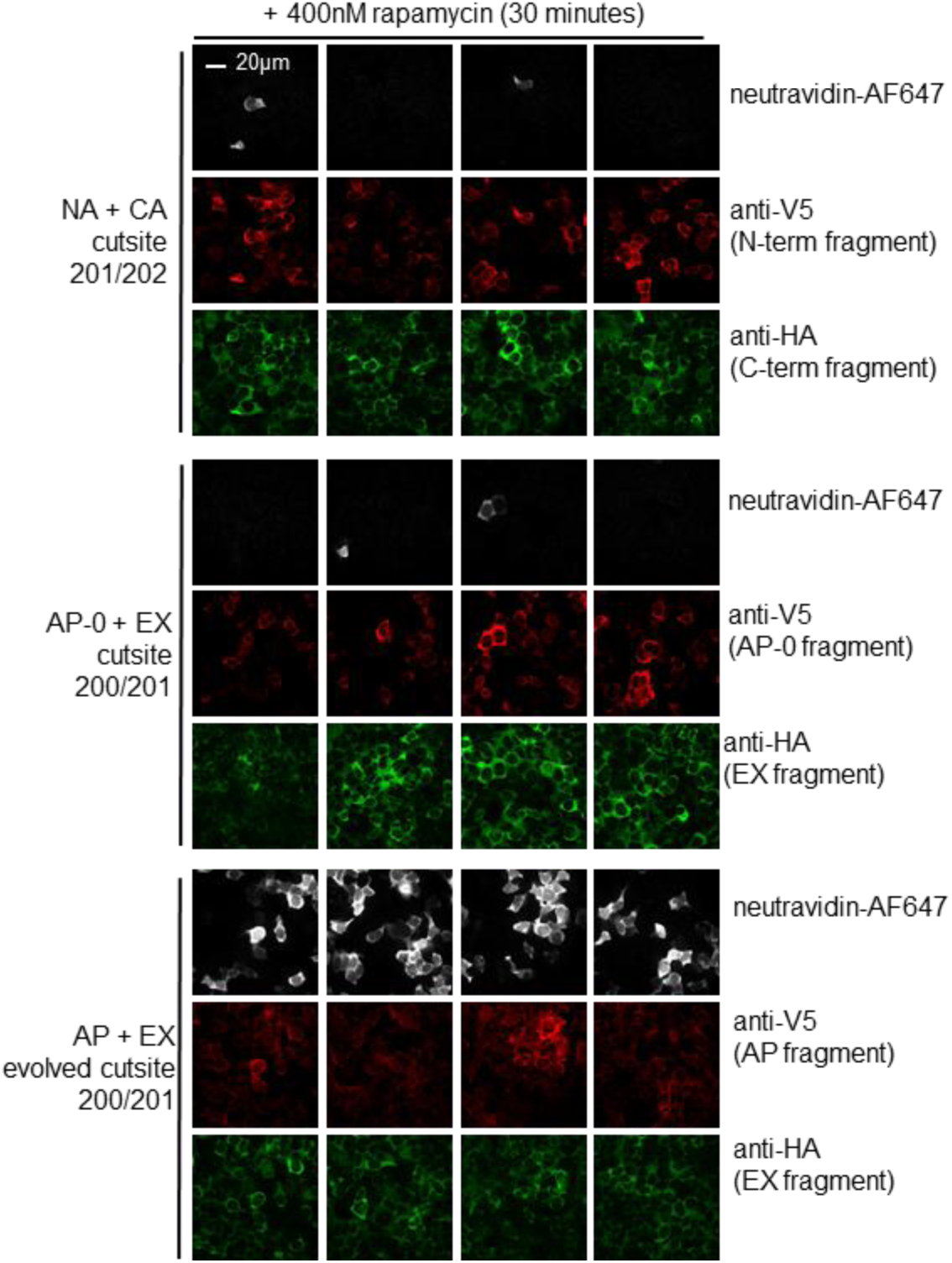
Comparing alternative split APEX2^57^, our original split APEX2, and evolved split APEX2. Comparison of sAPEX variants in the mammalian cytosol, with extent of biotinylation as the readout of peroxidase activity. See **Figure 3A** for depiction of split APEX2 protein fusions. Cut site 201/202 (NA and CA)^57^ was tested in tandem with our chosen cut site 200/201 (AP-0 + EX) as well as our final evolved version of AP and EX. The indicated N-and C-terminal variants of sAPEX were introduced by induction into HEK 293T cells. After protein production and heme supplementation, cells were incubated with BP for 30 minutes, a well as rapamycin to drive reconstitution. Cells were then labeled in the presence of H_2_O_2_ for 1 minute, fixed and stained with neutravidin-AlexaFluor647 to visualize peroxidase-catalyzed promiscuous biotinylation^2^. Four separate fields of view are shown per condition. Scale bar 20 µm. The experiment has been biologically replicated once.

**Figure S3.**
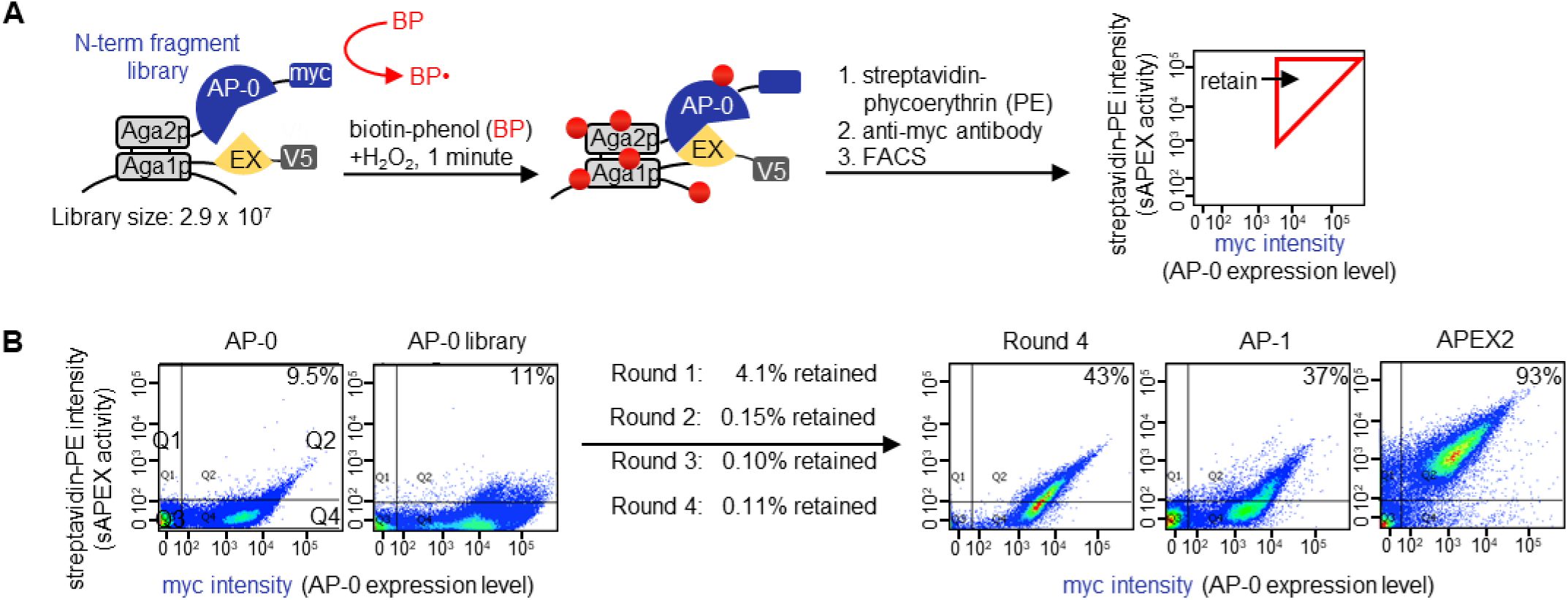
Generation 1 yeast display-based directed evolution. (**A**) Experimental setup. A library of N-terminal fragment AP-0 variants was displayed on the yeast surface via fusion to the mating protein Aga2p. A constant C-terminal fragment, EX (aa 201-250 of APEX2), is co-displayed via fusion to Aga1p. The yeast cell library is treated with biotin-phenol (BP) and H_2_O_2_ for 1 minute. Cells with active peroxidase self-biotinylate (red dots). Biotinylation sites are detected by staining with streptavidin-phycoerythrin (PE), and anti-myc antibody (followed by anti-Chicken-AlexaFluor647) is used to quantify AP-0 expression level. Using FACS, cells with a high PE/myc staining ratio were enriched. (**B**) FACS plots showing progress of selection. Cells were prepared and treated as in (**A**). AP-0 is the original N-terminal sAPEX fragment template. Round 4 is the population of yeast cells that remain after 4 rounds of FACS selection and re-amplification. AP-1 is the best clone from the Round 4 pool, whose mutations are shown in **Figure 2C**. “APEX2” is the original full-length APEX2. Percentage on the upper right corner of each FACS plot indicates the number cells in quadrant 2 divided by the total number of cells that are expressing N-terminal fragment (quadrant 2 + quadrant 4). The experiment has three biological replicates in total.

**Figure S4.**
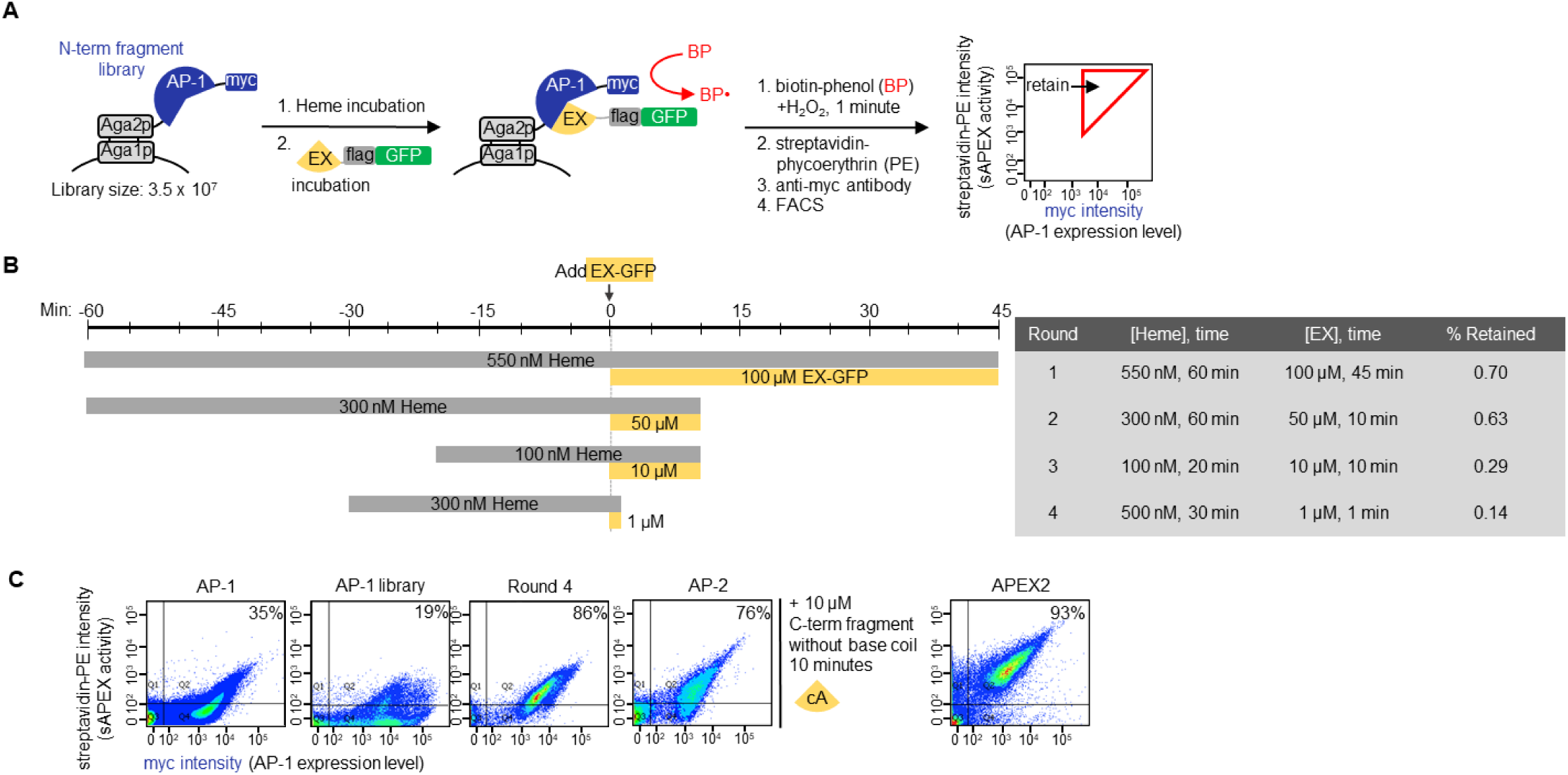
Generation 2 yeast display-based directed evolution. (**A**) Experimental setup. A library of N-terminal fragment AP-1 variants was displayed on the yeast surface via fusion to the mating protein Aga2p. The C-terminal fragment is added as a purified protein. BP labeling, streptavidin-phycoerythryin staining, and FACS are performed as in **Figures 2A and S3A**. (**B**) Specific labeling conditions used in each round of selection. (**C**) FACS analysis of indicated samples. Round 4 is the library population remaining after four rounds of selection. AP-2 is the best clone from 4 rounds of selection. Cells were incubated with 500 nM of heme for 30 minutes before addition of 10 μM EX-flag-GFP for 10 minutes followed by BP labeling, streptavidin staining, and FACS. Note that the fluorescence from GFP was not measured during FACS; GFP was included to increase the solubility of EX. Percentage on the upper right corner of each FACS plot indicates the number cells in quadrant 2 divided by the total number of cells that are expressing N-terminal fragment (quadrant 2 + quadrant 4). The experiment has two biological replicates.

**Figure S5.**
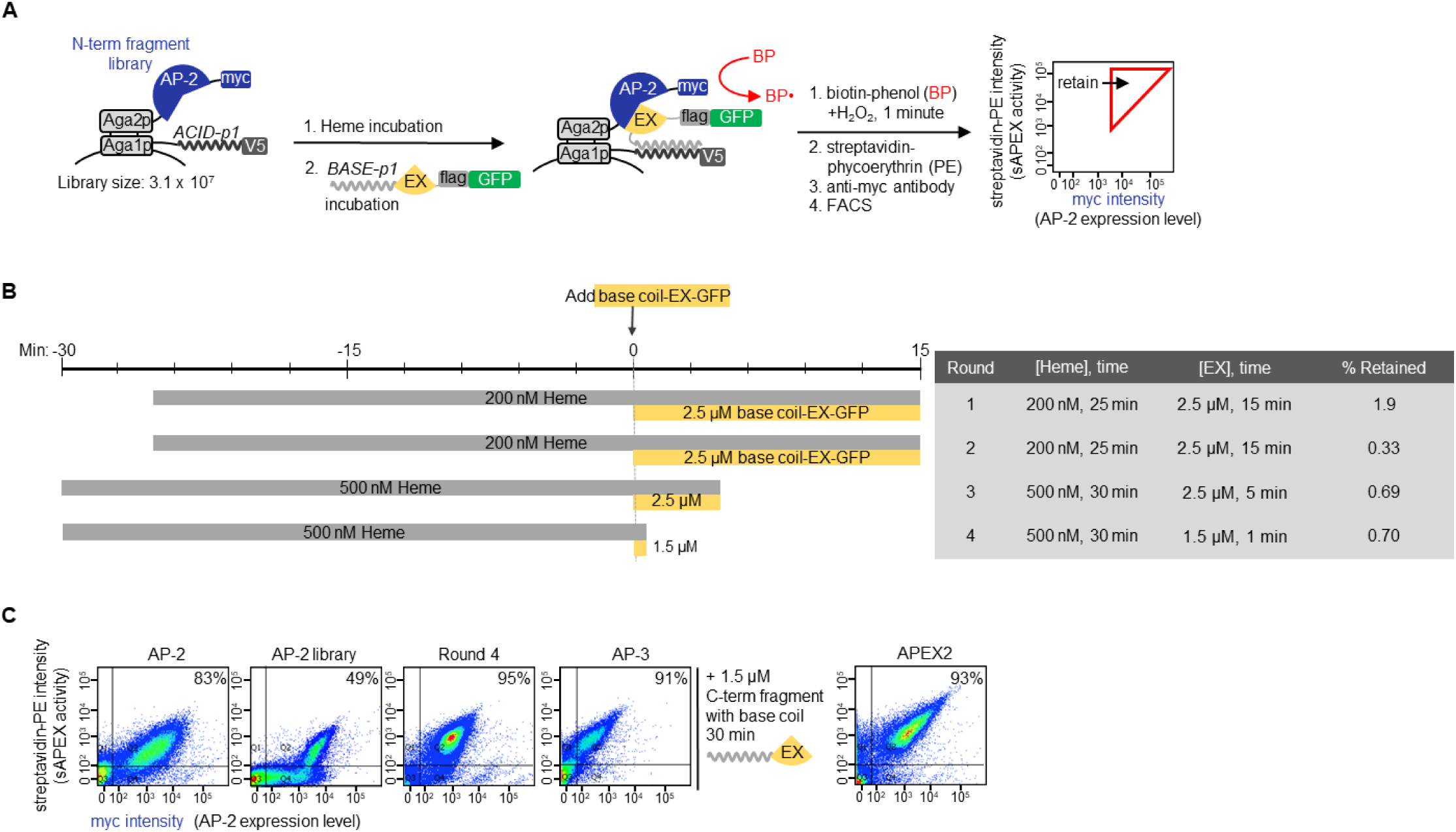
Generation 3 yeast display-based directed evolution. (**A**) Experimental setup. This is the same as the positive selection shown in **Figure 2A**. (**B**) Specific labeling conditions used in each round of selection. (**C**) FACS analysis of indicated samples. Labeling condition was 500 nM of heme for 30 minutes, then 1.5 µM base coil-EX-GFP protein for an additional 30 minutes, before BP labeling, streptavidin staining, and FACS. Note that the fluorescence from GFP was not measured during FACS; GFP was included to increase the solubility of EX. Percentage on the upper right corner of each FACS plot indicates the number of cells in quadrant 2 divided by the total number of cells that are expressing N-terminal fragment (quadrant 2 + quadrant 4). Two biological replicates.

**Figure S6.**
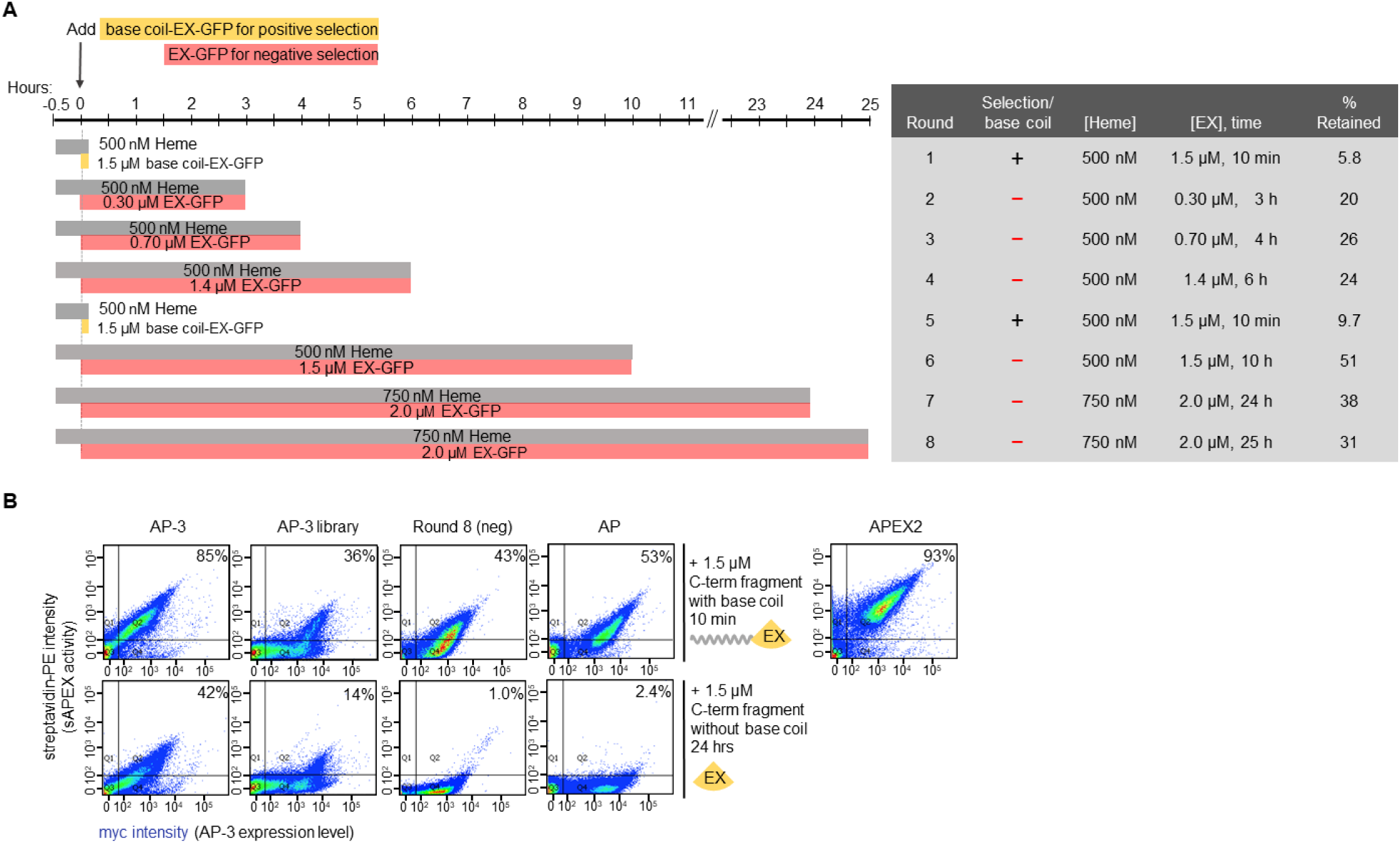
Generation 4 yeast display-based directed evolution. The experimental setup for this generation is shown in **Figure 2A**. (**A**) Specific labeling conditions used in each round of selection. (**B**) FACS analysis of indicated samples. Labeling condition was 500 nM of heme for 30 minutes, then 1.5 μM EX-GFP, with or without base coil, for 10 min or 24 hours, respectively. Next, BP labeling, streptavidin staining, and FACS were performed. Percentage on the upper right corner of each FACS plot indicates the number cells in quadrant 2 divided by the total number of cells that are expressing N-terminal fragment (quadrant 2 + quadrant 4). Two biological replicates.

**Figure S7.**
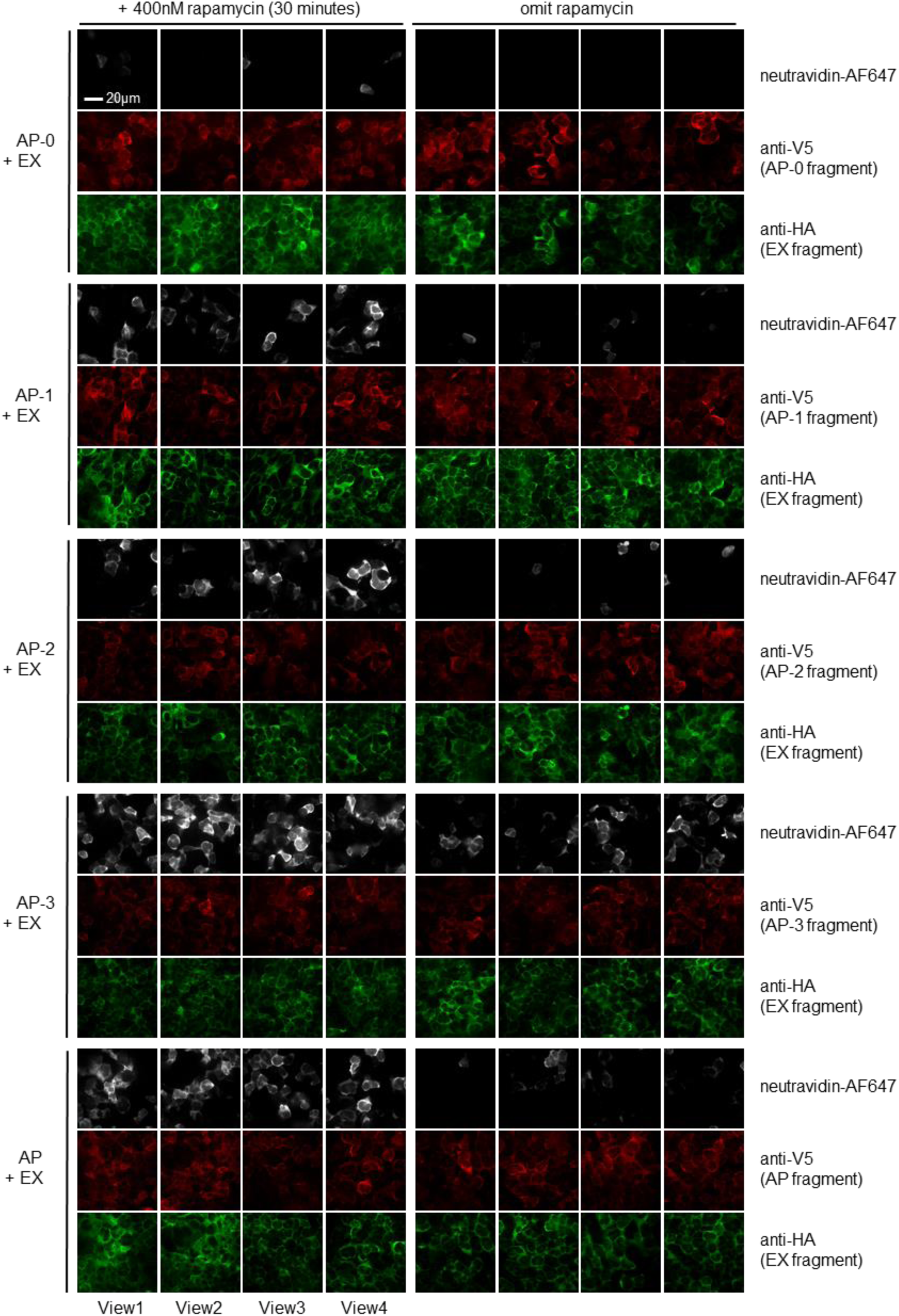
Comparison of sAPEX variants in the mammalian cytosol, with biotin-phenol labeling as readout of peroxidase activity. Images are from **Figure 3C**, but here anti-V5 and anti-HA channels are shown as well. Anti-V5 staining quantifies expression of the N-terminal fragment, AP, and anti-HA staining quantifies expression of the C-terminal fragment, EX. Scale bar, 20 µm.

**Figure S8.**
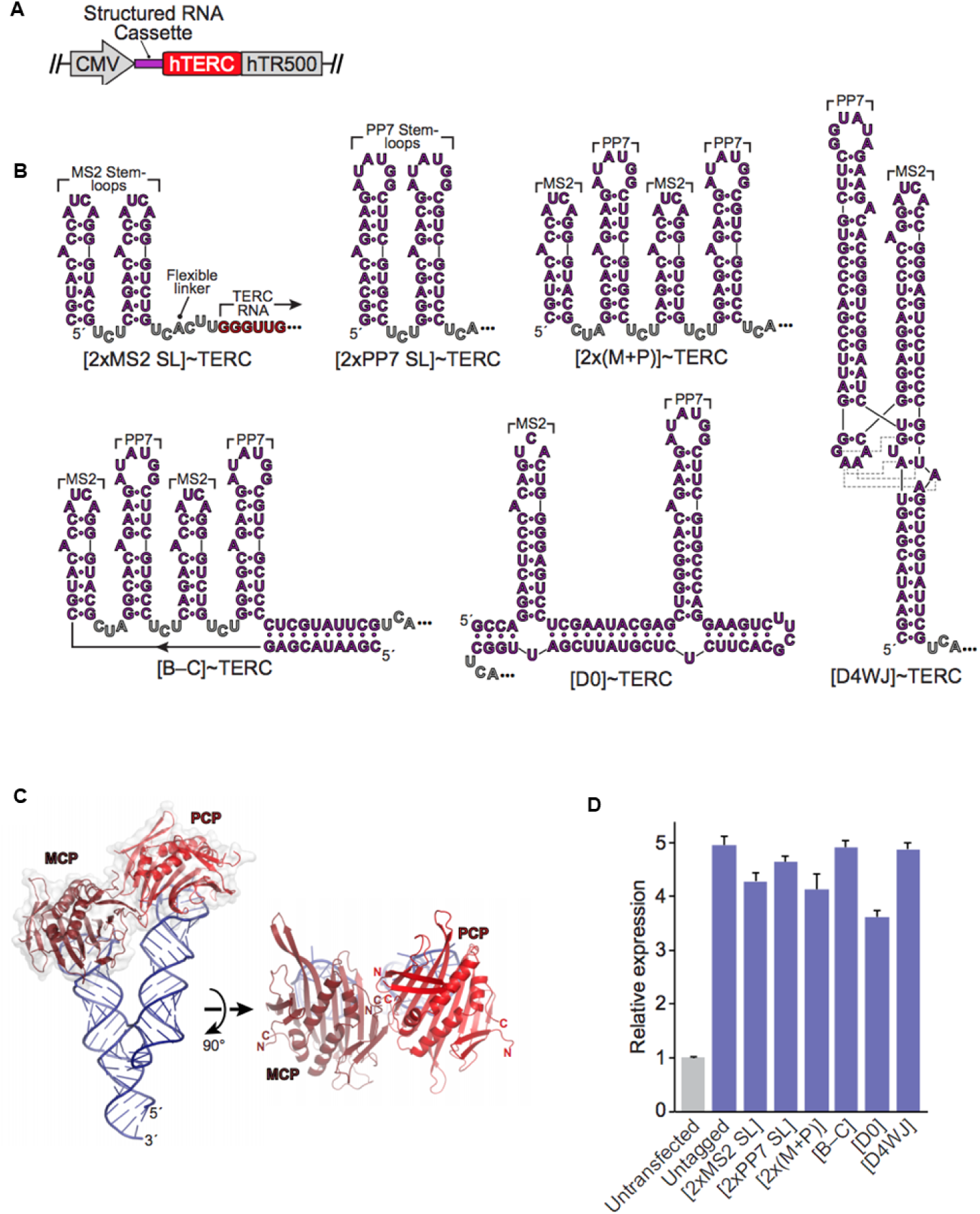
Noncoding RNA construct design. (**A**) General schematic of the tagged TERC RNA expression system. A cytomegalovirus promoter (CMV) drives expression of human telomerase RNA (hTERC, *red*) appended at its 5′ terminus with a structured cassette (*purple*). TERC 3′–end processing is mediated by the hTR500 block (*gray*), corresponding to the 500 bp of native genomic sequence downstream of the TERC 3′–terminus^71^. (**B**) Predicted secondary structures of the RNA cassettes tested. Linker residues denoted in gray; the 3′–terminal UCAUUU linker is present on all constructs. Red nucleotides indicate the 5′–end of the TERC ncRNA. Cassettes contain the following elements: [2xMS2 SL], two MS2 stem-loops; [2x PP7 SL], two PP7 stem-loops; [2x(M+P)], two pairs of alternating MS2 and PP7 stem loops; [B–C], a “bracketed cassette,” in which the [2x(M+P)] construct is enclosed within an additional helix; [D0], the “D0” RNA scaffold, previously demonstrated to co-localize MS2– and PP7–fusion proteins *in vivo*^94^; [D4WJ], a “docked 4-way junction” cassette, derived from “tecto-RNA” constructs designed to promote parallel co-proximation of two RNA helices^95^. Gray dotted lines denote tertiary contacts that mediate the interhelical docking interaction^96^. (**C**) Three-dimensional model of the D4WJ construct, bound to MCP and PCP (modeled on PDB IDs 2BU1, 2QUX, and 1GID). Orthogonal views are shown. Note the close proximity between the coat promoters’ N– and C–termini (“N” and “C”) near the protein–protein interface. Modeling was performed in COOT^97^; figure was generated using Pymol (Schrödinger, LLC). (**D**) Quantitative RT–PCR expression analysis of TERC constructs transiently expressed in HEK 293T cells. A common pair of primers that target the TERC RNA core was used to survey all constructs. Relative expression (scaled to untransfected control cells, which naturally express TERC) was quantified via the ΔΔC_T_ method, normalized to GAPDH. Data correspond to the mean of four technical replicates. Error bars: standard deviation.

**Figure S9.**
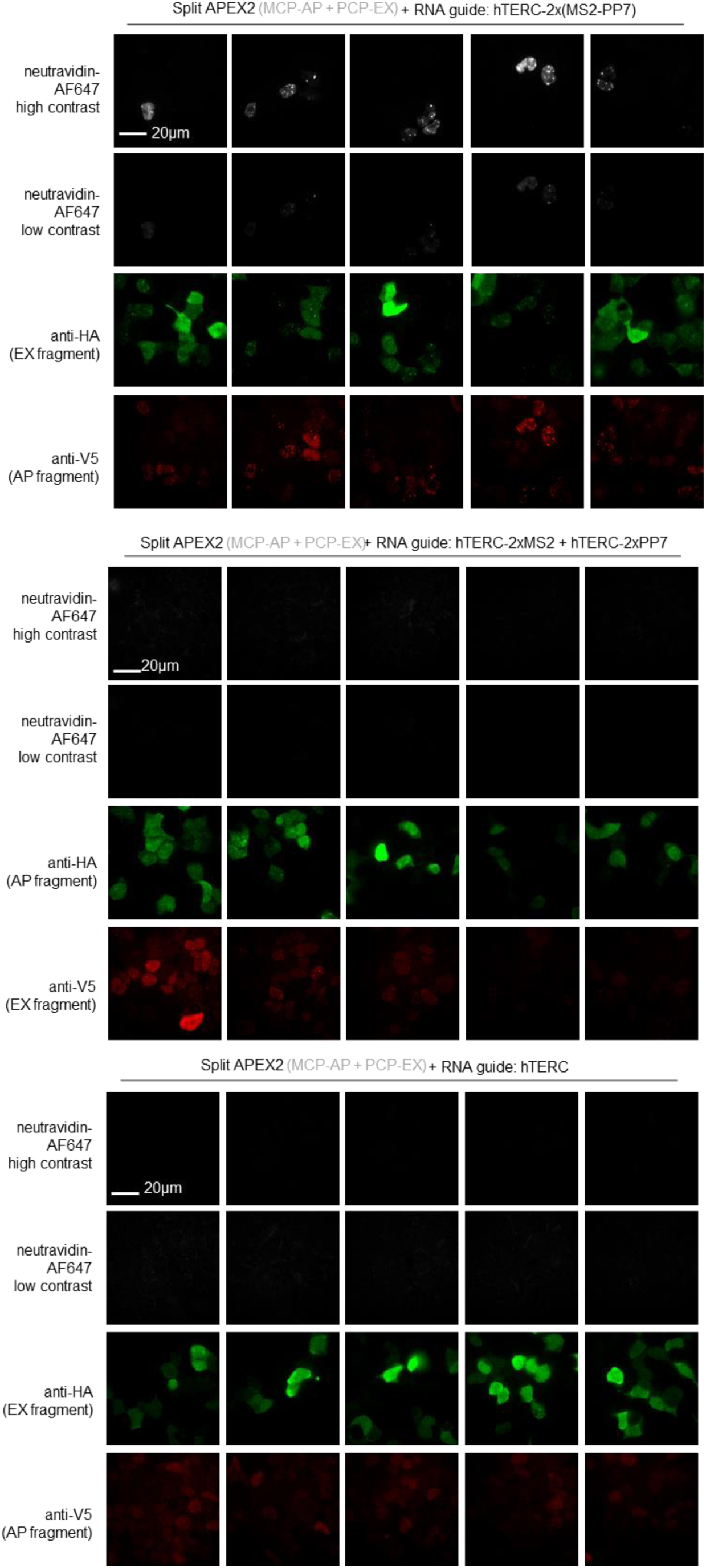
Additional fields of view of sAPEX reconstitution on RNA binding sites. Additional fields of view of **Figure 4C.** Clonal HEK 293T cells stably expressing MCP-AP were transfected with PCP-EX and different hTERC RNA guides. Scale bar, 20 µm.

**Figure S10.**
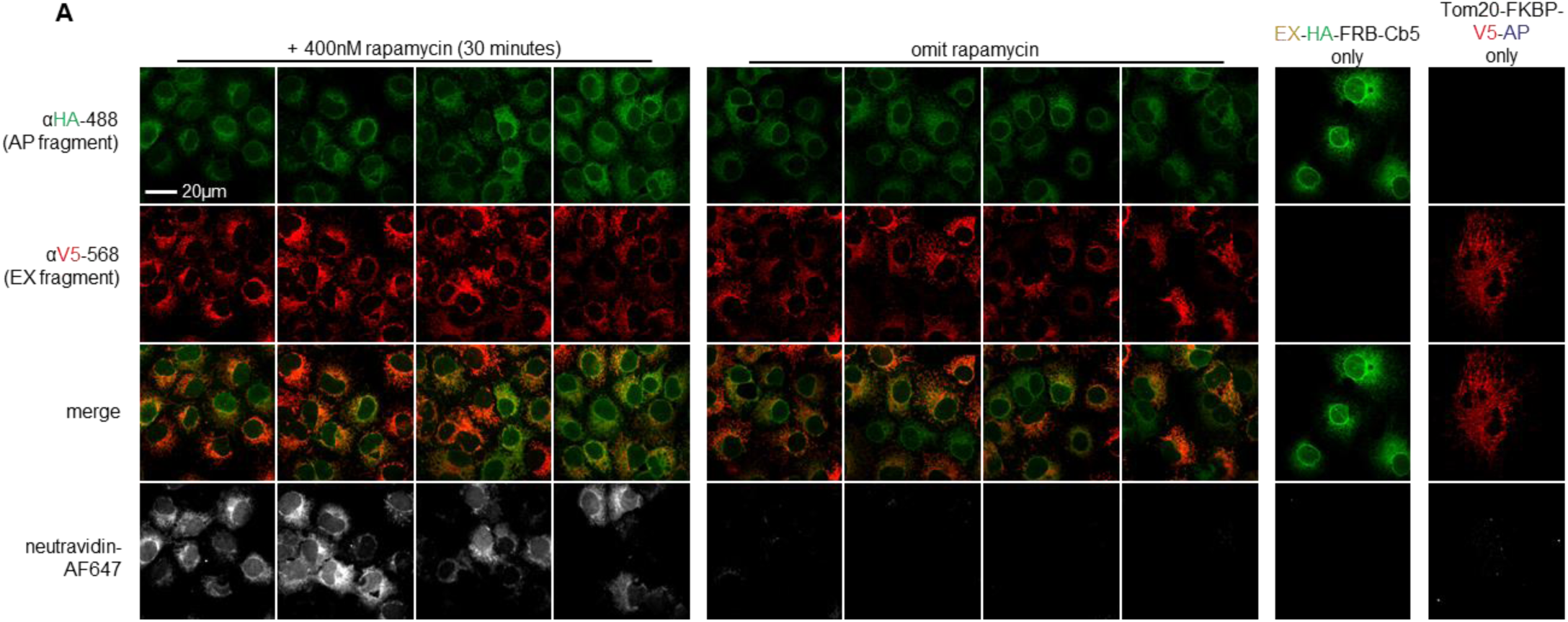
Examining sAPEX mito-ER targeting and morphology. (**A**) Additional fields of view for mito-ER experiment shown in **Figure 4F.**Controls are also shown with each sAPEX fragment alone (right columns). Scale bar, 20 µm. Note that because BP labeling was performed on living cells, biotinylated proteins can diffuse away from the site of labeling, leading to a diffuse neutravidin staining pattern^2,3^.

**Figure S10.**
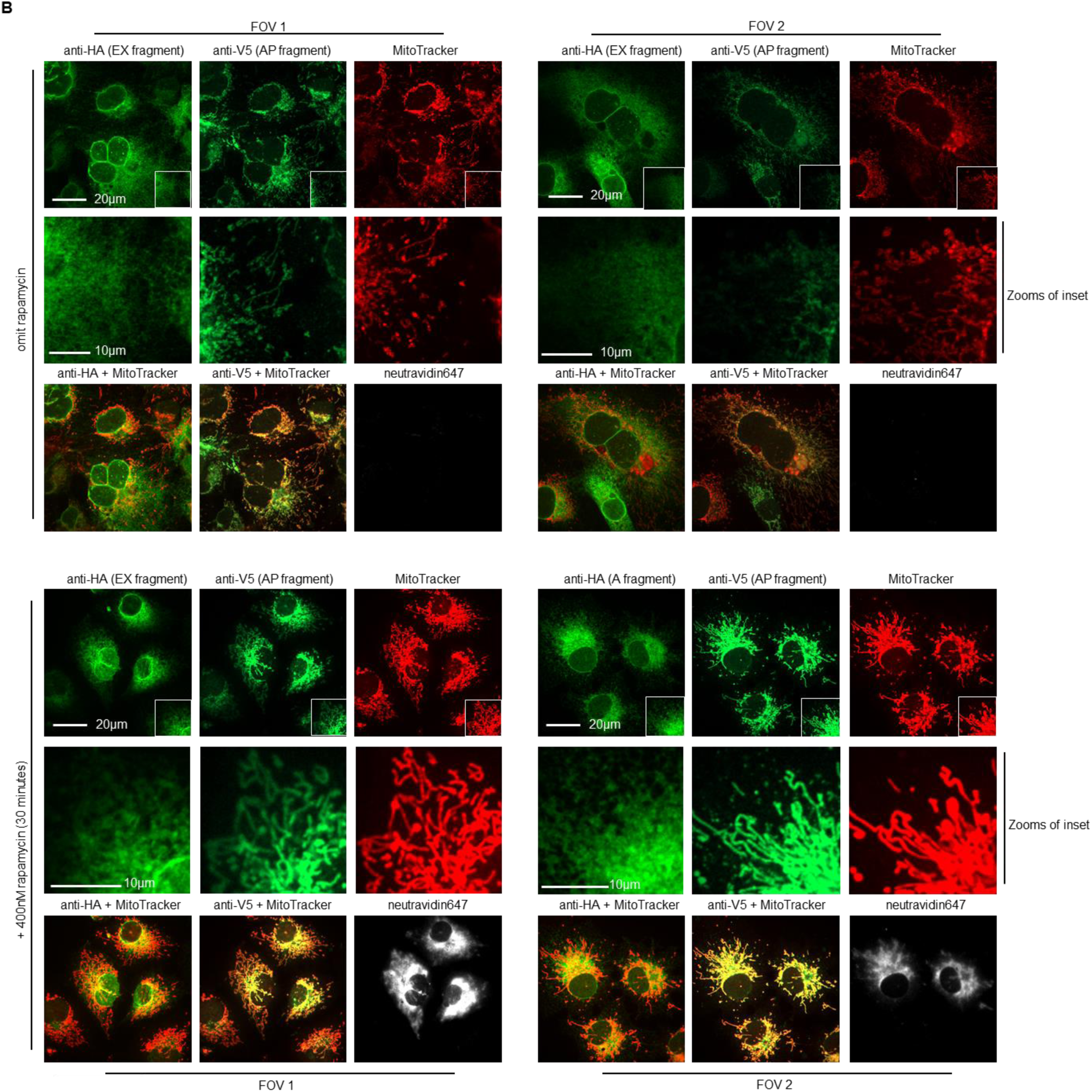
Examining sAPEX mito-ER targeting and morphology. (**B)**With matched conditions to **Figure 4F**, the morphology of the mitochondrial was visualized during confocal microscopy using MitoTracker Red. Scale bar, 20 µm unless otherwise indicated. The experiment has been biologically replicated twice.

**Figure S10.**
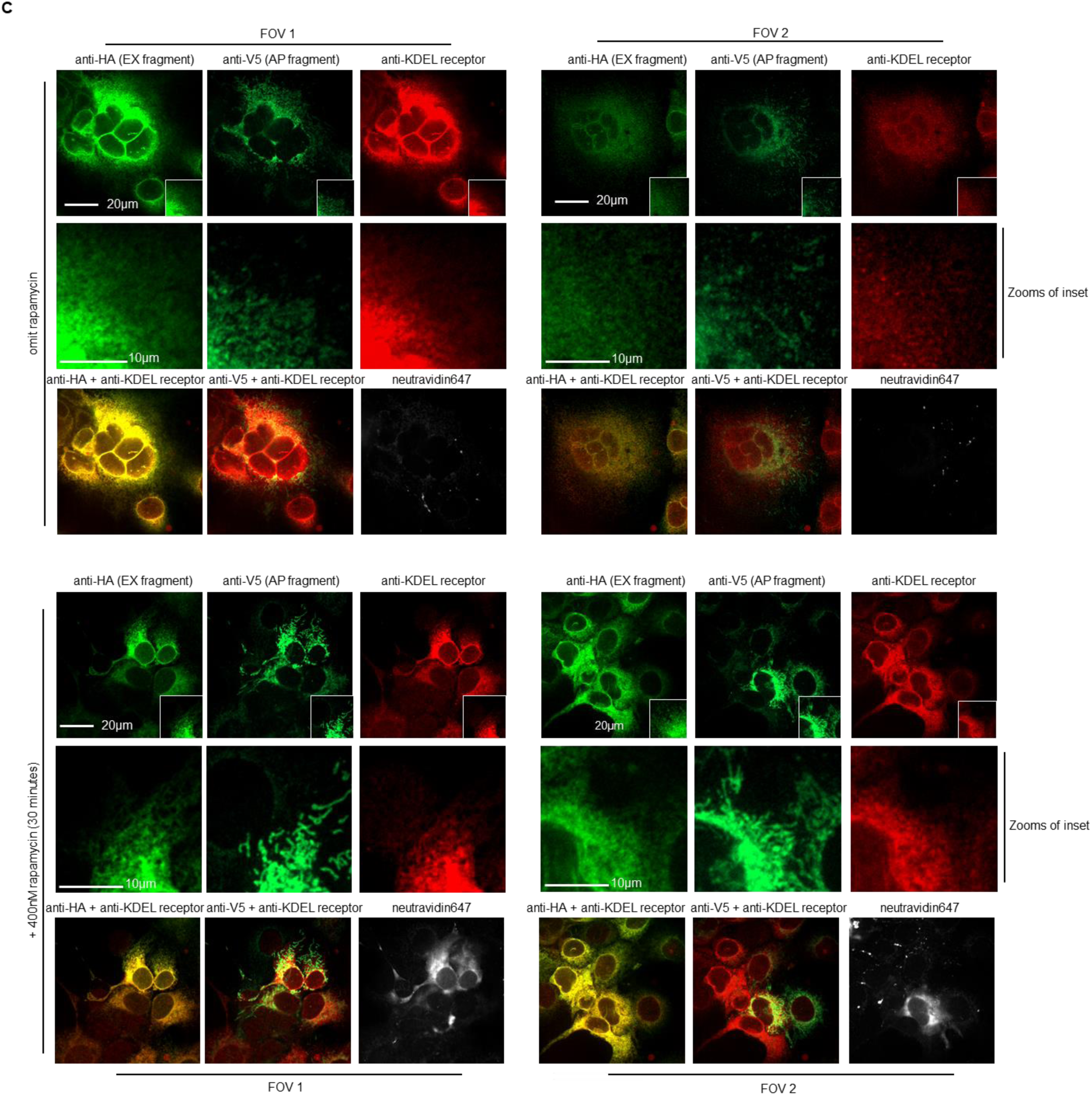
Examining sAPEX mito-ER targeting and morphology. (**C)**Same as **S10B**, except the morphology of the ER was visualized during confocal microscopy using an antibody against the KDEL-receptor. Scale bar, 20 µm unless otherwise indicated. The experiment has been biologically replicated once.

